# Distinct gene expression dynamics in developing and regenerating limbs

**DOI:** 10.1101/2021.06.14.448408

**Authors:** Chiara Sinigaglia, Alba Almazán, Marie Sémon, Benjamin Gillet, Sandrine Hughes, Eric Edsinger, Michalis Averof, Mathilde Paris

## Abstract

Regenerating animals have the ability to reproduce organs that were originally generated in the embryo and subsequently lost due to injury. Understanding whether the process of regeneration mirrors development is an open question in most regenerative species. Here we take a transcriptomics approach to examine to what extent leg regeneration shows the same temporal patterns of gene expression as leg development in the embryo, in the crustacean *Parhyale hawaiensis*. We find that leg development in the embryo shows stereotypic temporal patterns of gene expression. In contrast, global patterns of gene expression during leg regeneration show a high degree of variation, related to the physiology of individual animals. A major driver of this variation is the molting cycle. After dissecting the transcriptional signals of individual physiology from regeneration, we obtain temporal signals that mark distinct phases of leg regeneration. Comparing the transcriptional dynamics of development and regeneration we find that, although both processes use largely the same genes, the temporal patterns in which these gene sets are deployed are different and cannot be systematically aligned.

**HIGHLIGHTS:**

- Single-limb data on transcriptional dynamics of leg development and regeneration
- Developing embryonic legs show stereotypic transcriptional profiles
- Regenerating leg transcriptomes show a high degree on individual variation
- Regenerating leg transcriptomes are influenced by adult physiology, especially molting
- Regenerating leg transcriptomes reveal distinct phases of leg regeneration
- Leg development and regeneration use overlapping sets of genes in different temporal patterns

## INTRODUCTION

Many animals have the capacity to regenerate body parts that have been lost after a severe injury. In some cases regeneration produces faithful replicas of the lost organs, which are indistinguishable from those originally developed in the embryo. This similarity in the *outcome* of development and regeneration suggests that the *processes* generating these structures could also be similar, i.e. that regeneration could mirror embryonic development. The fact that both take place within the same organism, relying on the same genome, makes it easy to envisage that the same gene regulatory networks could be used in both cases.

In spite of this evident connection, however, there are important ways in which development and regeneration are likely to differ. Development is a stereotypic process, unfolding from a defined starting point, in the stable and well-provisioned environment of the fertilised egg. In contrast, regeneration starts with an injury whose extent and timing are unpredictable. Regeneration unfolds in the context of adult physiology, influenced by the nutritional status of the animal, exposure to microbes, as well as circadian, seasonal and/or other hormonal cycles. For instance, in arthropods, the molting cycle profoundly affects the physiology of the individual and imposes physical constraints on the growth of regenerating structures (Charmantier-Daures and Vernet 2004).

In the adult body, the cellular context in which regeneration takes place differs from the embryo: different pools of progenitors are available compared with development, and differentiated cell types such as immune cells and neurons (which are not yet formed at the onset of limb development) are known to play key roles in supporting regeneration (e.g. Kyritsis et al. 2012, Godwin et al. 2013, Sinigaglia and Averof 2019).

Significant differences can also be seen in the scales in which development and regeneration unfold. Embryonic organ primordia are usually hundreds of micrometres to millimetres scale, while adult regenerating organs can be orders of magnitude larger (see Brockes and Kumar 2005). Such differences in scale are likely to have an impact on the mechanisms that coordinate cell behaviour and cell fate across developing tissues, such as diffusion-based morphogen gradients and long range cell-cell communication. Such differences may also apply to the temporal scales in which development and regeneration unfold.

Over several decades, numerous studies have compared aspects of development and regeneration in efforts to address these questions. Several studies provide evidence of significant similarities in the patterning mechanisms that operate during development and regeneration, in axolotl limbs (e.g. Muneoka and Bryant 1982, Roensch et al. 2013; reviewed in Nacu and Tanaka 2011) and in other systems (Aztekin et al. 2021, Czarkwiani et al. 2021), while others point to significant differences (Fan et al. 2012, Bosch et al. 2010, Warner et al. 2019, Khan and Crawford 2020).

The crustacean *Parhyale hawaiensis* presents an excellent system for exploring the relationships between embryonic and regenerative processes, for several reasons. First, *Parhyale* are able to regenerate their legs with high fidelity; regenerated legs are indistinguishable from the original, unharmed legs (A. Almazan, C. Cevrim, M. Paris and M. Averof, unpublished). Second, *Parhyale* are direct developers (do not undergo metamorphosis), so the adult legs directly derive from the legs developing in the embryo. Third, although adult legs are larger than embryonic legs, the leg primordia in the developing embryo and regenerating adult develop on similar scales. The primordia measure in the order of 100 micrometres and consist of a few hundred cells (Alwes et al. 2016, Wolff et al. 2018). The temporal scales of leg development and regeneration are also similar, spanning 4-5 days at 26°C from primordium/blastema formation to fully patterned leg (Browne et al. 2005, Alwes et al. 2016, Wolff et al. 2018). These shared attributes render *Parhyale*’s developing and regenerating legs an attractive system in which to test the possible ways in which the mechanisms of regeneration could mirror those of development.

On a practical level, the developing and regenerating legs of *Parhyale* are easily accessible at all stages, and have previously been studied by continuous live imaging at cellular resolution (Alwes et al. 2016, Wolff et al. 2018). The *Parhyale* genome has been sequenced and annotated (Kao et al. 2016, see also Supplementary Materials) providing a valuable resource for transcriptomic studies. Finally, transgenic reporters and tools to manipulate gene function have been developed in this species, opening the way to functional genetic studies in the future (reviewed in Paris et al. 2021).

To compare gene usage during development and regeneration on a genome-wide scale we performed a large-scale RNAseq on single legs, covering the time course of each process: from early limb buds/freshly amputated legs to fully patterned legs (96-192 hours post fertilization and 0-150 hours post amputation, respectively). Our sampling focused on homologous portions of the distal leg in developing and regenerating legs. Our working hypothesis was that some phases of leg regeneration, such as wound closure, are likely to be specific to regeneration, while others like patterning, morphogenesis, growth and/or cell differentiation could share significant similarities. Our goals were: 1) to compare expression dynamics on a global scale to determine whether specific phases of leg regeneration are most similar -in terms of gene expression-to specific phases of leg development, 2) to identify the sets of genes associated with each of these phases and determine whether a significant fraction of these co-expressed genes are shared between development and regeneration, and 3) to determine whether, overall, these sets of co-expressed genes are deployed in the same temporal order in the embryo and in the regenerating adult leg.

Our expectation was that we should be able to recover common temporal patterns of gene expression underpinning the embryonic and regenerative time courses, consistent with the idea that some aspects of leg regeneration re-deploy mechanisms used for leg embryonic development. Alternatively, a failure to detect common temporal patterns of gene expression would suggest that development and regeneration are coordinated by distinct mechanisms.

## RESULTS AND DISCUSSION

### Transcriptional profiling of leg development reveals stereotypic developmental profiles

To investigate transcriptional dynamics and assess individual variation in developing embryonic legs we performed RNAseq on individual T4 legs, sampled every 6 hours from 96 to 192 hours post fertilization (hpf). This sampling captures the entire time course of leg development from a young limb bud stage to fully patterned and differentiated legs (Figure 1a, Browne et al. 2005). In order to account for the progressive regionalization of the primordium, we collected a complete series of samples consisting of the entire T4 leg (F series) and, when possible, the distal portion of the T4 leg, corresponding to the distal 1/3 of the leg from 120 to 138 hpf (AB series), and to the carpus, propodus and dactylus from 144 to 192 hpf (A series). The complete and the distal leg samples were collected in pairs, from contralateral T4 legs of the same embryos. We collected at least two replicates for each timepoint and sample type, yielding a total of 70 samples.

**Figure 1:**
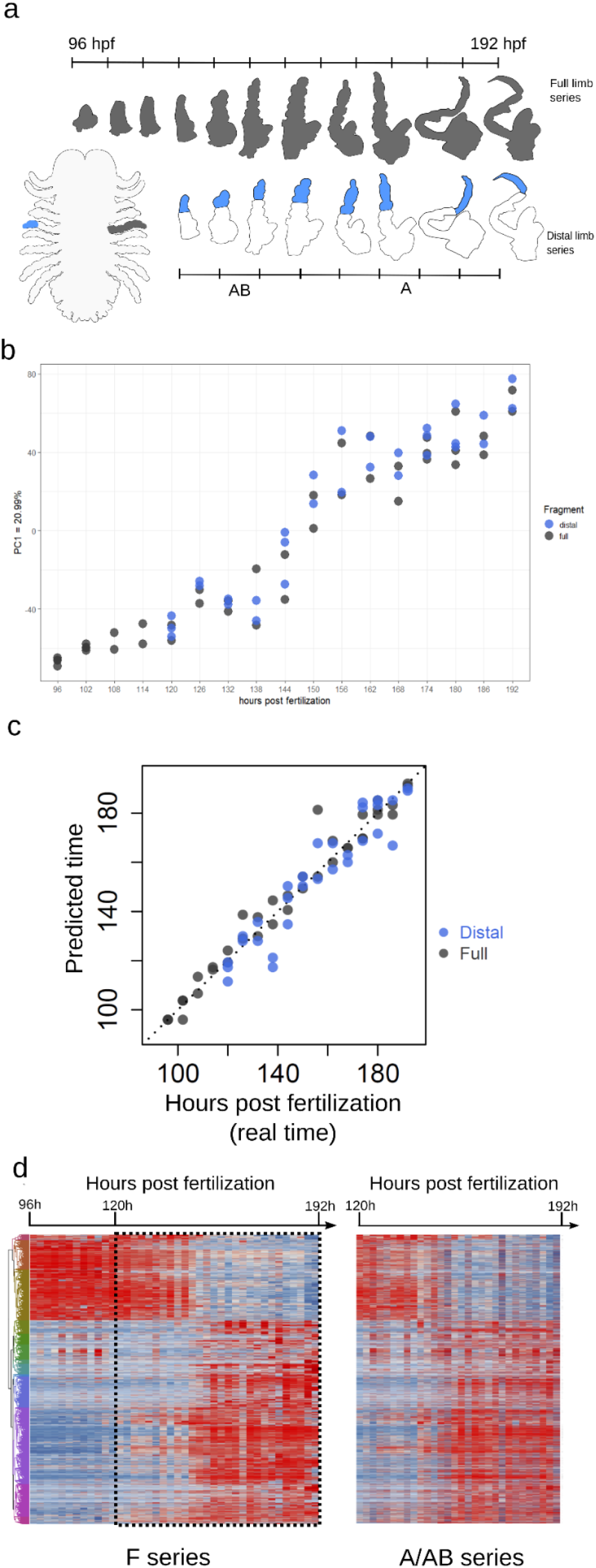
Transcriptional profiling of leg development reveals stereotypic developmental dynamics. (**a**) General morphology of a *Parhyale* embryo and description of the experimental plan for building the timeline of leg embryogenesis. The dissection plans of the F and A/AB series are highlighted in grey and in blue, respectively. (**b**) Principal component analysis (PCA) of the F and A/AB series. Samples separate along PC1 (y axis) mainly according to developmental stage (hour post fertilization, x axis). The proportion of the variance explained by the first components of the correspondence analysis is indicated. (**c**) The stage of the samples from the A/AB series was predicted using RAPToR on a reference built from the samples from the F series (y axis) and correlates well with time after fertilization (x axis). (**d**) Heatmap of expression dynamics in the F and A/AB series for genes that vary during sampled time-line (8196 genes differentially expressed). The time-window in the F series that corresponds to the time-window of the A/AB series is highlighted with a dashed rectangle.

Principal component analysis (PCA) on the complete RNAseq dataset showed that the main axis of gene expression variation, explaining 52% of the variance, strongly correlates with developmental time in both the F (full) and the A/AB (distal) leg samples (Figure 1b). To probe the strength of the temporal signal in those data, we applied RAPToR, a method that allows to predict the developmental stage of a sample from its gene expression profile, relative to a reference time series (https://github.com/LBMC/RAPToR). We built a reference using the F leg samples and used this to estimate the stage of each of our samples. The predictions of the model match the real developmental age of each sample accurately, not only for the F samples (which were used to train the model) but also for the A/AB leg samples (Figure 1c). These results indicate that the temporal dynamics of gene expression in developing legs are highly reproducible, and that the temporal dynamics captured in the F and A/AB series are highly coherent with each other.

Comparing further the transcriptional dynamics of F and A/AB samples during embryogenesis, we found 7963 and 1354 genes to be differentially expressed during embryogenesis in F and A/AB samples respectively (DESeq2, padj < 0.001, see Supplementary materials), with an overlap of 1121 genes (the complete lists of annotated genes are provided in the Supplementary Materials). We attribute the higher number of differentially expressed genes in the F samples to the fact that this dataset spans a longer developmental period and includes additional tissues. The tissue dissections to collect the A/AB samples were also more challenging, possibly contributing to lower sample quality (Supplementary materials). In spite of these differences, we noticed a high similarity in gene expression dynamics between these two datasets (Figure 1d). Overall, this analysis shows that the temporal signal of the distal part of the leg (A/AB) is largely recapitulated in the full leg series (F).

### Transcriptional dynamics of leg regeneration are masked by individual variation

We performed a similar series of RNAseq experiments to investigate the temporal dynamics of gene expression during the course of regeneration in adult T4 legs, amputated at the distal end of the carpus. Previous studies have shown that the cellular activity associated with regeneration occurs within 200 microns from the amputation site (Konstantinides and Averof 2014, Alwes et al. 2016), which in these experiments corresponds approximately to the distal half of the carpus. Samples were collected every 12 hours, from the moment of amputation until 120 hours post amputation (hpa), when the legs appear to be fully patterned (Alwes et al. 2016). To ensure that we have sampled patterned and differentiated legs, we also collected samples at the onset of expression of a late distal leg marker (the *Distal*^*DsRed*^ exon trap; Kontarakis et al. 2011, Konstantinides and Averof 2014) and after the first molt following regeneration.

From each of these limbs we collected two samples: one consisting of the distal-most end of the limb stump (the carpus, including the blastema and newly regenerating structures; series A) and one from a more proximal podomere (the distal part of the merus; series B) (Figure 2a). The A samples capture the entire part of the limb that participates actively in regeneration (Alwes et al. 2016), while the B samples serve as controls, intended to capture transcriptional variations associated with the physiological status of each individual (e.g. molting stage, nutritional state, etc.). Overall, we collected 120 samples from 37 individuals (paired samples were collected from the left and right T4 legs in 23 individuals), spanning 13 time points, yielding a total of 60 A and 60 B samples (Figure 2a).

**Figure 2:**
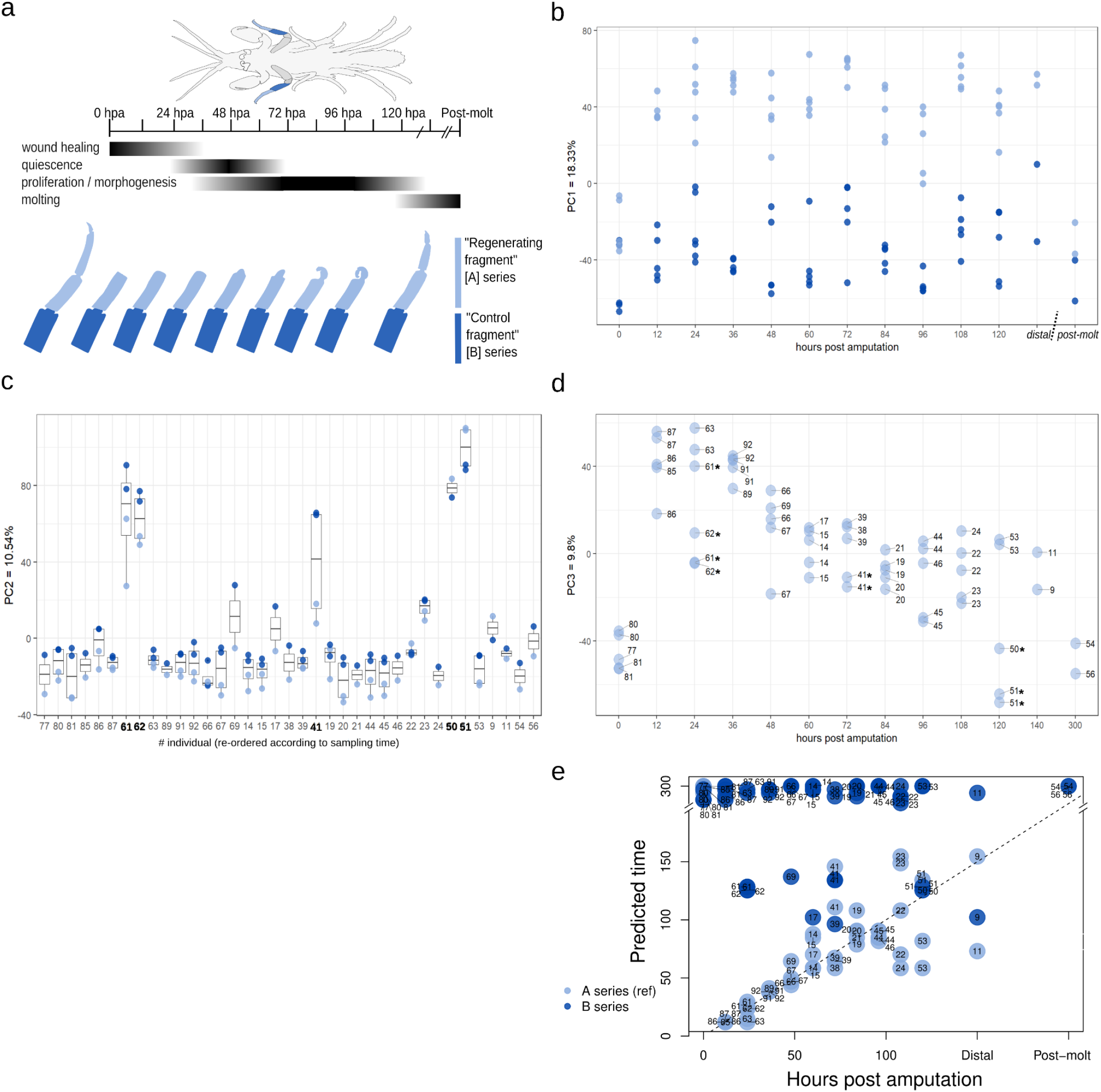
Transcriptional dynamics of leg regeneration are masked by individual variation. (**a**) General morphology of a *Parhyale* adult and description of the experimental strategy. For each limb, we sampled a regenerating fragment (A series), and a more proximal control fragment (B series), which are highlighted in light blue and in dark blue, respectively. The timing of the different phases of regeneration, as established by live imaging, is given (**b-d**) PCA of the A and B series. **(b)** Samples separate along PC1 (y axis) mainly according to whether the sample is regenerating, not according to time after amputation (x axis). (**c**) Samples separate along PC2 according to the individual animal they were collected from (x axis). (**d**) “A” samples broadly separate according to time after amputation (x axis). The proportion of the variance explained by the first components of the correspondence analysis is indicated. (**e**) The stage of the samples from the B series is very poorly predicted using a RAPToR-reference built on the samples from the A series.

Principal component analysis of the A and B series reveals several distinct sources of variation in these data. PC1 captures the difference between the A (regenerating) and B (control) samples, with the notable exception of the 0 hpa (unamputated) and the post-molt A samples, which groups together with the B series (Figure 2b). This grouping suggests that tissues undergoing active regeneration are transcriptionally distinct from the non-regenerating samples. There is no evidence of a temporal signal in PC1.

PC2 also appears to lack a temporal signal, revealing instead marked differences between the groups of samples collected from different individuals (particularly in individuals marked in bold, Figure 2c). We return to the source of this individual variation in the next section.

A temporal signal, mirroring the time course of regeneration, is clearly visible in the A samples in PC3 (Figure 2d). On this axis, the transcriptional profile of pre-amputation samples (0 hpa) matches the profile of samples collected at the end of the regenerative time course, consistent with our expectation that regeneration is complete by ∼120 hpa and after the following molt.

Although the temporal dynamics of regeneration account for only a small part of the total variation in these samples (PC3, explaining 10% of the variance) we have several reasons to think that these data are of high quality and are largely capturing biological variation, rather than noise. First, PC1, PC2 and PC3 are capturing real sources of biological variation: regenerating tip versus non-regenerating proximal tissue in PC1 (Figure 2b), physiological differences related to the molting cycle in PC2 (see next section and Figure 3b), and the temporal dynamics of regeneration in PC3 (Figure 2d). Second, we are confident that the inter-individual variation recovered in PC2 reflects real biological differences, rather than sampling noise, as we find a much higher correlation of gene expression in samples collected from the same individual than among different individuals, and the values found in A and B samples are significantly correlated within individuals (Figure 2c and Supplementary materials). Third, when we apply RAPToR to the regenerating samples, using a reference timeline built with all the A and B samples, we find that the gene expression data are able to predict the stage of regeneration of each A sample quite accurately (Figure 2e). In contrast, the RAPToR model makes very poor predictions on the B samples, which are not expected to carry a regenerative signal.

**Figure 3:**
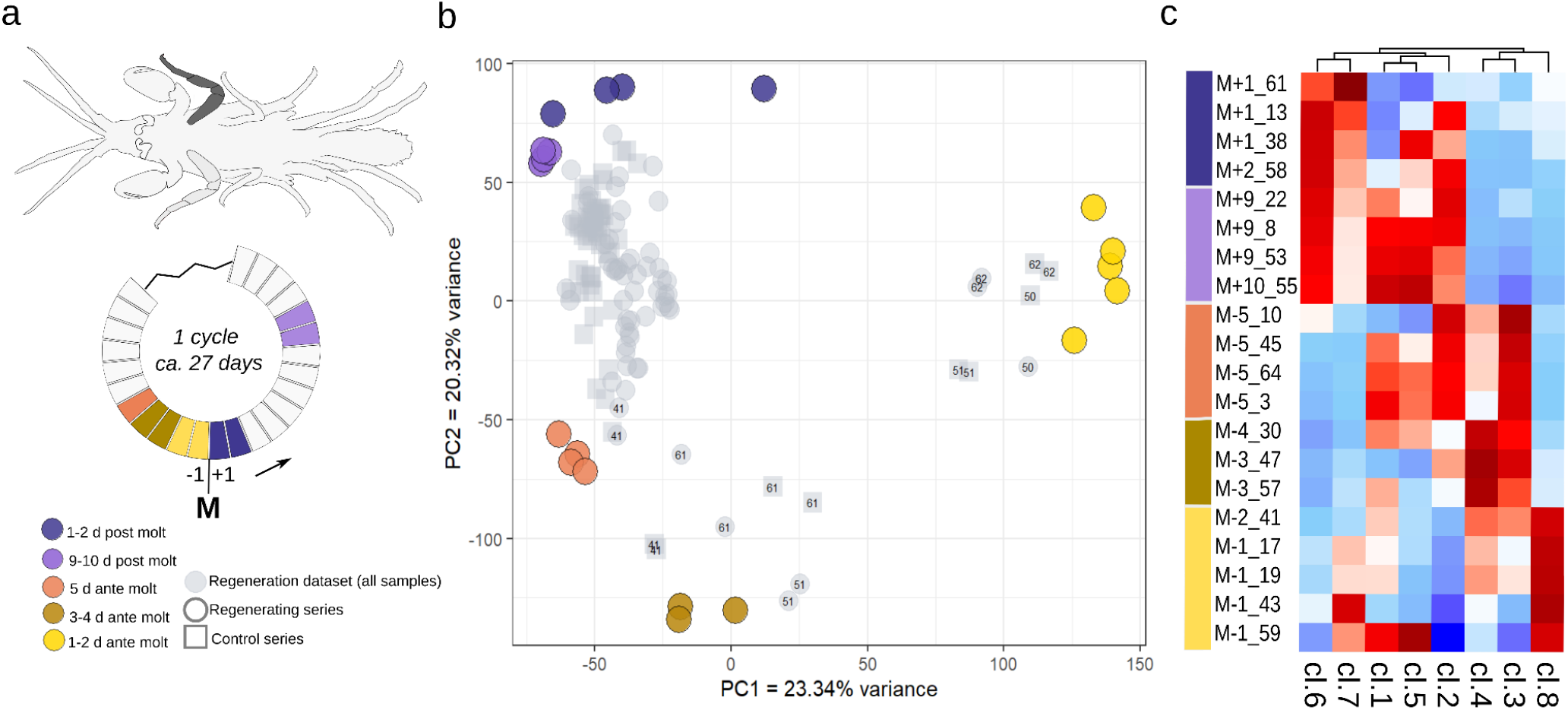
The molting cycle has a strong impact on the transcriptional profile of adult limbs. (**a**) One entire T4 limb (in grey) was sampled from animals at different stages of the molting cycle: sampling strategy is indicated on the wheel below, which depicts an average molting cycle. (**b**) PCA of the 20 molting samples (large circles). Samples separate on PC1 and PC2 according to their molting stage. The proportion of the variance explained by the first components of the correspondence analysis is indicated. Values for the regeneration dataset (in grey) were projected onto the PCA. The outlier regenerating samples detected in Figure 2c are closest to the samples about to molt (yellow and brown circles), the identifier of the individual is reported. (**c**) Fuzzy c-means clustering of the expression values from the molting dataset. Centroid values of each of the 8 clusters are color-coded from -2 (dark blue) to +2 (dark red).

### The impact of the molting cycle on the transcriptional profile of adult limbs

We hypothesise that the observed ‘individual signal’ (seen in PC2, Figure 2c) might be linked to the physiological state of each animal, as it is shared by all the samples collected from one individual. Since molting is a major physiological variable in adult crustaceans, we decided to directly test the impact that the molting cycle has on the leg transcriptome.

Selecting animals of the same age/size as in the regeneration RNAseq experiments (see Methods), we monitored the molting status for 66 animals over two successive molts; we observed that this cohort molted with a mean period of 27 days (SD 7.2). We then collected entire T4 legs from 20 of these animals at different stages of the molting cycle: 5 days, 3-4 days and 1-2 days prior to molting, and 1-2 days and 9-10 days after molting (3 to 5 replicates per condition, Figure 3a) and performed RNAseq on the samples. Principal component analysis on these 20 samples shows that different stages of the molt cycle are well separated on PC1 and PC2, representing almost half of total variation (large circles on Figure 3b). This analysis reveals sharp changes in the transcriptional profile of legs, taking place between 5 days pre-molt and the time of molting (orange, brown and yellow points in Figure 3b), compared with stable transcriptional profiles found in the intermolt period, starting from 1-2 days after the molt (blue and purple points in Figure 3b).

On the same principal components, describing the molting cycle, we projected the expression data of our regeneration time series, in order to assess the molting status of these samples and to probe the impact that the molting cycle might have on inter-individual variation. We observed a very good correspondence between the position of the regeneration samples on this projection (revealing their molting status) and the inter-individual variation revealed previously in the regeneration time series (PC2 in Figure 2c): most of the samples fell in the intermolt phase, except those that were previously marked as showing significant individual variation (marked by a star in Figures 2 and 3), which were associated with the near-molt stages (Figure 3b).

Applying a soft clustering approach (mFuzz, see Methods) on the molting cycle dataset, we defined eight distinct sets of co-expressed genes, which corroborate the idea that samples collected during the intermolt have similar transcriptional profiles (blue and purple phases in Figure 3c) and that samples collected shortly before molting show significant changes in gene expression (orange, brown and yellow phases in Figure 3c).

Our analysis therefore confirms our initial hypothesis that molting status has a strong transcriptional influence on the regenerating leg transcriptomes; in particular, the imminence of molting deeply modifies the transcriptional state of an adult leg, to an extent that overshadows the signal from leg regeneration (Figures 2c-d).

### Disentangling the transcriptional signals of physiology and regeneration

To investigate the transcriptional dynamics driven by the regenerative process independently of the physiological/molting status of each animal, we developed a Bayesian modeling approach (using JAGS; Plummer 2003) to dissect the contributions of regeneration (R) and the individual’s physiology (Cont) on overall transcriptome measured in A and B samples (grey circles in Figure 4a). Based on the results presented earlier, we assumed that the variation due to an individual’s physiology would be shared by all the samples collected in each individual (including A and B and paired left/right samples). In contrast, the variation in gene expression driven by the regenerative process should be specific to the A samples. Previous observations suggested that individual limbs can regenerate at different speeds (Alwes et al. 2016, and C. Sinigaglia, M. Lebel and M. Averof, unpublished observations); we therefore modeled the regenerative signal separately in each A sample, even when they were collected from the left and right T4 legs of the same individual.

**Figure 4:**
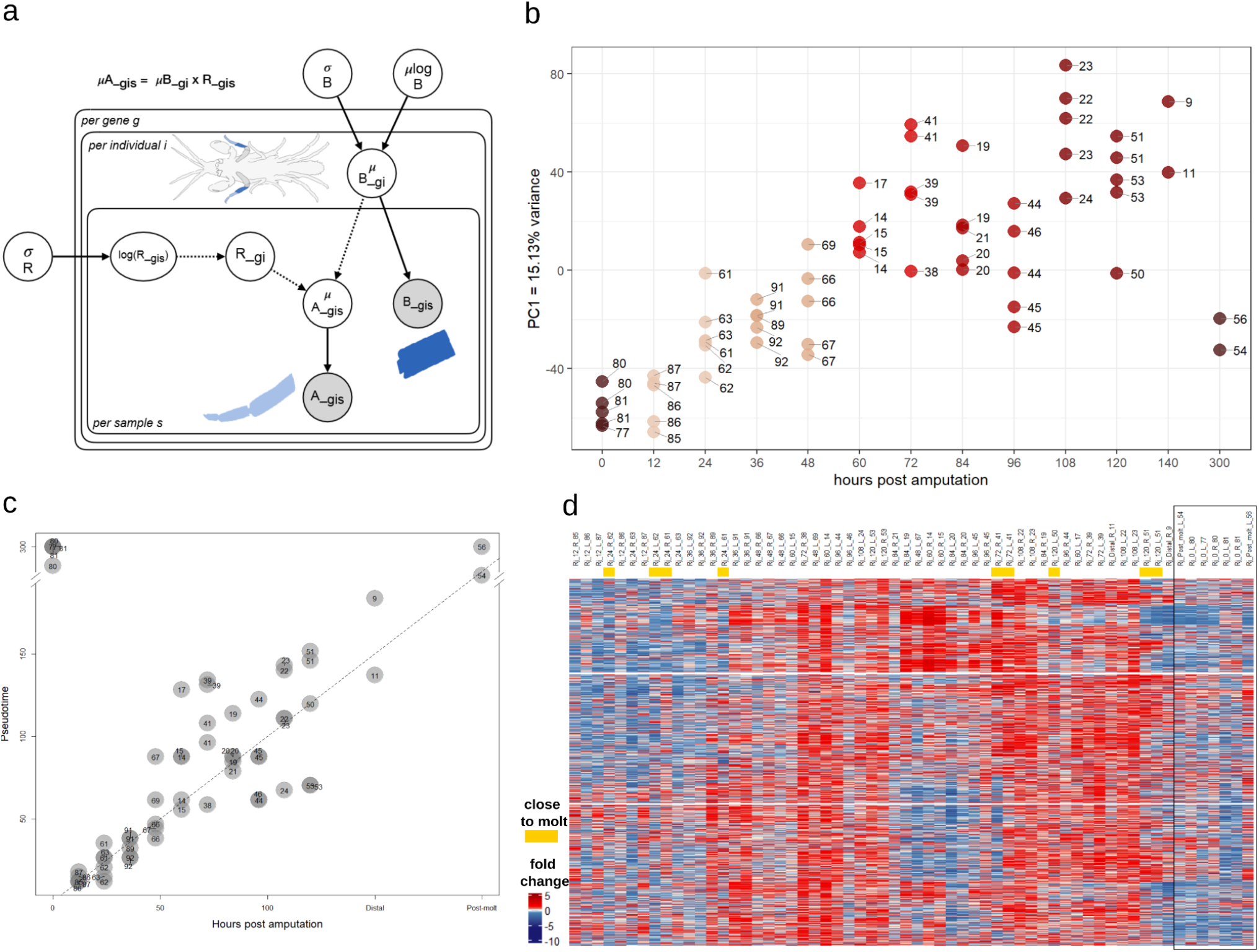
Modeling of the regenerative signal. (**a**) Directed acyclic graph illustrating the model used to extract R values from the raw counts of A and B series. (**b**) Principal component analysis of the R values: Samples separate along PC1 (y axis) mainly according to regeneration stage (hour post amputation, x axis). Artificially assigned time-from-amputation were given to the distal and post-molt samples (150 hpa and 300 hpa, respectively). (**c**) Correlation between sampling time (x axis) and pseudotime (estimated with RAPToR, y axis) for the R values. For predicting pseudotime, the pre-amputation samples were considered as the terminal point of the regenerative sequence, and have been arbitrarily assigned a sampling time of 300 hpa. (**d**) Heatmap depicting the expression dynamics of the 12000 genes contributing the most to the regenerative/temporal signal in the R samples (PC1 in **b**). Samples taken from the individuals identified to be close to molt are annotated in yellow. Samples are ordered according to pseudotime (estimated in **c**) which places both non-amputated and post molt samples at the end of the timeline.

The regenerative signal R was modelled as an enrichment factor, such that the measured expression values for each gene in each A sample are equal to R multiplied by the individual’s control/physiological signal (taking into account sampling errors/variation). An R value of 1 conveys that there is no difference in expression between A and B samples, R>1 means that a gene is upregulated in the regenerating sample compared with B, and R<1 that a gene is downregulated. Further details on the modeling approach are given in the Methods and Supplementary Materials. Following this approach, we take R values to represent the effect that regeneration has on each gene and each sample along the regenerative time course.

To test whether this modelling approach is successful in recovering the expression dynamics of regeneration from the overall transcriptional variation, we performed a principal component analysis on the R series. We find that the major axes of variance now follow the time course of regeneration (PC1 shown in Figure 4b, PC2 in Supplementary materials). Using the R values we are also able to predict the phase of regeneration of each limb more accurately than with the raw count values, using RAPToR (Figure 4c). As seen earlier, the samples collected at the end of the regenerative time course have a similar profile to the pre-amputation samples (0 hpa).

We observe that the regenerative phase of each sample is predicted most accurately in early phases of regeneration (0-36 hpa) and least accurately in later phases (48-120 hpa), suggesting that transcriptional profiles are less accurately predicted by the time elapsed from amputation at later phases. This is consistent with what we have observed in live imaging experiments, in which wound closure and a period of quiescence occur reliably in the first 1-2 days of regeneration, but the onset of later events (cell proliferation and morphogenesis) varies (Alwes et al. 2016). Given these variations in the onset of some phases, the predictions made by RAPToR can help us to place the regenerating samples on a common temporal scale (‘pseudotime’) based on their regenerative transcriptomes.

To visualise the transcriptional dynamics captured in the R signal, we plotted R over pseudotime, focusing on the genes that contribute most significantly to PC1 in Figure 4b (Figure 4d). Despite our efforts to exclude the physiological/molting signal from R, we observe some residual differences in the samples collected shortly before molting (yellow highlights in Figure 4d).

### Distinct transcriptional dynamics in developing and regenerating legs

Having captured the transcriptional profiles of leg development and leg regeneration, we turn our attention to the goal of comparing these profiles, to determine whether some of the dynamics of leg regeneration could mirror those of leg development. First, we performed a principal component analysis combining both the regenerating samples (R values) and the embryonic leg samples (data from the F series). In the latter, raw counts were transformed into fold changes to render them comparable with R values (Bayesian modeling, see Methods); we refer to these transformed data as the E series.

This analysis shows that the embryogenesis dataset harbors more variation since the two major axes of variation (PC1 and PC2, explaining together 42% of the total variance) capture very clearly the transcriptional dynamics of leg development, but very poorly the dynamics of leg regeneration (Figure 5a). There is no obvious alignment between the axes of variation of the embryonic and the regenerating samples.

**Figure 5:**
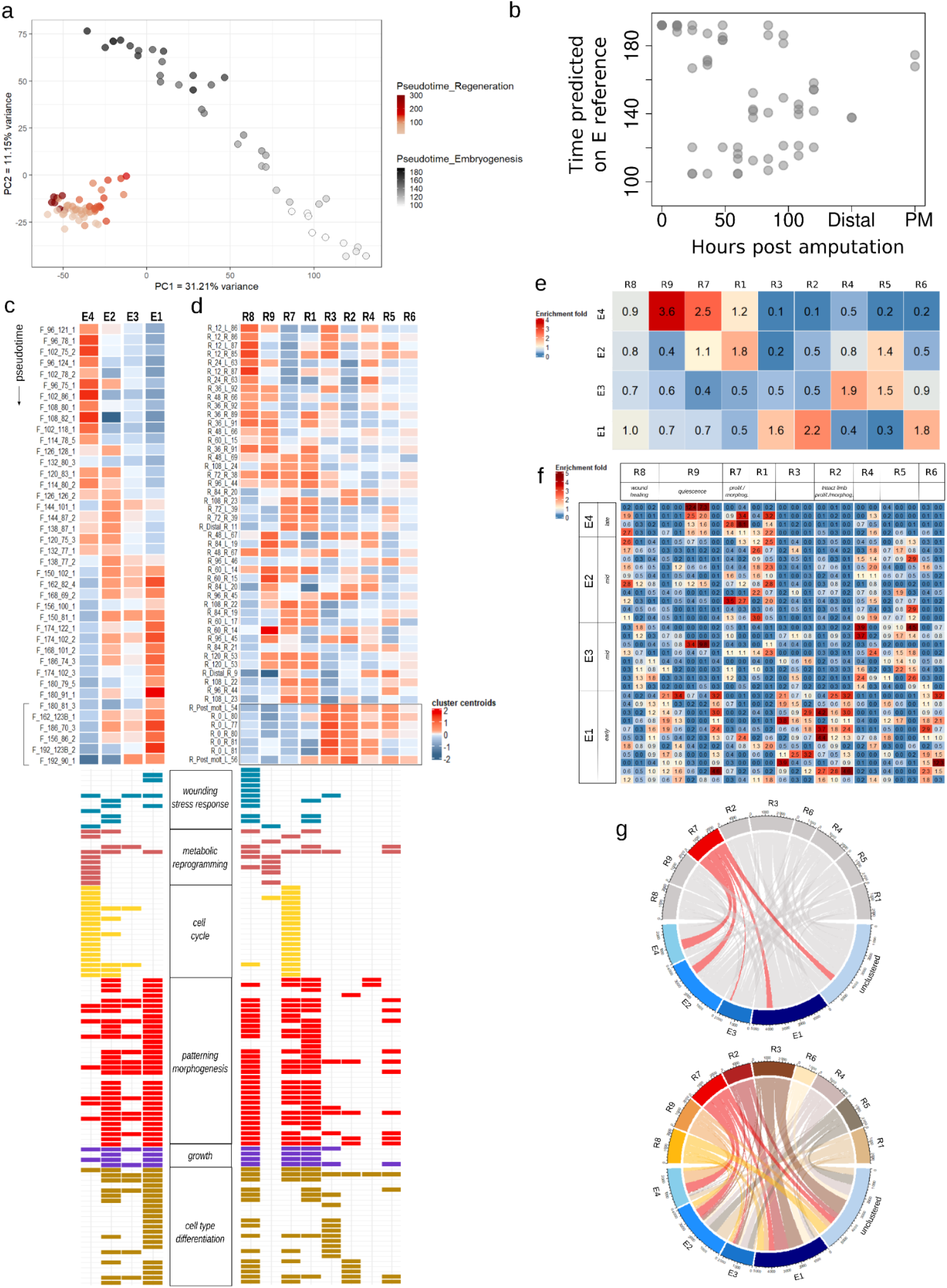
Comparison of developmental and regenerative transcriptional dynamics. (**a**) Principal component analysis of development and regeneration datasets (E and R values). Samples have been color-coded according to the predicted time (R in reds, E in greys). PC1 separates the two datasets, both PC1 and PC2 correlate with embryonic development. (**b**) RAPToR alignment of R sample timeline (sampling time) to the embryonic reference time. (**c**) Fuzzy c-means clustering of the expression values and summary of the GO enrichment analysis, for the E values. The 4 clusters have been reordered on the x axis according to their temporal dynamics. Cluster sizes are the following: E4 - 3107, E3 - 3444, E1 - 6641, E2 - 5950. Top: heatmap of cluster centroids; samples have been re-ordered according to pseudotime (y axis). Bottom: Summary of enriched GO terms, trimmed down with Revigo and color coded according to relevant categories. (**d**) Fuzzy c-means clustering of the expression values and summary of the GO enrichment analysis, for the R dataset. The 9 clusters have been reordered on the x axis according to their temporal dynamics. Cluster sizes are: R8 - 2331, R9 - 2153, R7 - 2497, R1 - 2401 genes, R2 - 1966 genes, R3 - 2874, R4 - 2301, R5 - 2159, R6 - 1413. Top: heatmap colors represent the cluster centroid values. Samples have been re-ordered according to predicted time from Figure 4c (y axis), which places the pre-amputation and post-molt samples at the end of the timeline. Bottom: summary of enriched GO terms (same as in c).(**e**) Overlap of embryonic and regenerative clusters, expressed as enrichment folds relative to random clusters of same size. Clusters have been reordered as in c and d. (**f**) Overlap of higher resolution embryonic and regenerative clusters (25 for R and 33 for E). Correspondence between low and high resolution clusters has been determined with clustree. Clusters have been reordered following the temporal dynamics seen in c and d. (**g**) Graphical depiction of the genes shared between regenerative and embryonic clusters (R9, red-brown palette vs E4, in blue). Top: chord diagram highlighting the R7 cluster (in red), corresponding to the late regenerative phase of proliferation and morphogenesis. Bottom: match between all regenerative and embryonic clusters. A fraction of genes from the regeneration clusters (> 5000) are not clustered in the embryonic dataset.

Next, we tried to temporally align these datasets using RAPToR, by building a reference time series based on the embryonic leg (E) data and applying this to make temporal prediction on the regenerating leg (R) data. Pre-amputation and post-molt samples are consistently assigned to the latest stages of the leg development series, as expected of fully differentiated leg tissues (Figure 5b). The other samples are inconsistently placed on the developmental series, some matching early and some late phases of development, suggesting that there is no straightforward way to relate the phases of leg regeneration with those of leg embryonic development.

Global expression profiles, probed by PCA and RAPToR, could be dominated by specific groups of genes that show strong differential expression (e.g. terminal differentiation genes), concealing expression dynamics that might occur on a smaller scale, but include genes that are relevant for comparing the mechanisms of development and regeneration (e.g. genes involved in patterning, the control of cell proliferation and morphogenesis). To dissect the temporal dynamics of specific sets of genes we turned to clustering approaches, which identify sets of genes that share similar expression profiles. For this, we applied a soft clustering implemented through Mfuzz to identify groups of co-expressed genes separately within the E and the R datasets, which classified genes into 4 and 9 co-expression clusters, respectively, for developing and regenerating legs (an alternative, finer classification yielded 25 and 33 clusters, respectively; see Methods for tuning of cluster numbers). The numbers of genes assigned to each cluster are indicated in the legend of Figure 4. Most of these clusters appear to be associated with a specific phase of leg development/regeneration. In the embryonic leg data, genes in cluster E4 are expressed predominantly in the early phases of leg development, genes in E2 in mid phases, and genes in E3 and E1 in the late phases (Figure 5c). In the regenerating leg data, genes in cluster R8 are specifically expressed in the early phases of regeneration, R9 in the early-mid phases, R7 and R1 in the mid-late phases, and R3 and R2 are associated with differentiated limbs (pre- and post-regeneration); clusters R4-R6 do not carry a strong temporal signal (Figure 5d).

Having defined these clusters of co-expressed genes, we compared the sets of genes contained in each cluster to determine the degree of overlap between the developmental and the regenerative gene clusters. The most significant overlap is observed between early developmental cluster E4 and the early-mid regenerative clusters R9 and R7 (Figure 5e,g). A good overlap is also observed between the mid developmental cluster E2 and the late regenerative cluster R1, and between late developmental and regenerative clusters associated with differentiated limbs (Figure 5e). These overlaps are also conserved when comparing clusters of smaller size (Figure 5f). In spite of these similarities, we do not observe a coherent pattern suggesting that similar gene clusters could be deployed during development and regeneration in a similar temporal sequence.

### Comparing the temporal dynamics of specific functional categories of genes in leg development and regeneration

To associate these gene clusters with biological functions, we performed a GO enrichment analysis on each cluster. We then grouped many of those GO terms in categories that describe processes of development and regeneration (see Methods; Figures 5c-d bottom). Embryonic leg clusters E4, E2-3 and E1 are enriched in genes associated with cell proliferation, patterning/morphogenesis and cell differentiation, respectively (Figure 5c), consistent with the processes that take place during the corresponding phases of leg development (Wolff et al. 2018). In leg regeneration, this analysis also yields results that agree with what we have learned about *Parhyale* leg regeneration from live imaging studies (Alwes et al. 2016). Live imaging of leg regeneration in *Parhyale* shows that the first day post amputation is dominated by wounding responses. This is reflected in the enrichment of the early R8 cluster in genes associated with wounding and stress responses (turquoise GO terms in Figures 5c-d). Next, comes a period of quiescence of variable duration, in which little cellular activity is visible under the microscope. During this phase we have noticed that cell nucleoli become enlarged (Supplementary material), a sign of increased ribosome biogenesis and metabolic activity (Tiku and Antebi 2018). Interestingly, the R9 cluster that becomes upregulated 36-48 hpa is highly enriched in genes associated with ribosome biogenesis and metabolism (dark red GO terms in Figure 5d).

The quiescence phase is followed by a period of active cell proliferation and morphogenesis. Among the co-expression clusters active during this phase, from 48 to 120 hpa, cell proliferation appears to be specifically associated with cluster R7 (yellow GO terms in Figures 5c-d) and clusters R7 and R1 are strongly enriched in genes associated with patterning, morphogenesis and growth (red and purple GO terms). Clusters that are predominantly associated with pre-and post-regenerative phases (R3 and R2) are enriched in GO terms connected with cell differentiation (brown GO terms). Overall, we observe that these GO annotations are enriched in clusters of both datasets, but they are not distributed in the same way across embryonic and regenerative co-expression clusters. Notably, leg regeneration appears to have unique early phases associated with wound response and metabolic reprogramming, followed by a phase that is characterised by overlapping profiles for cell proliferation and patterning and morphogenesis. In contrast, embryonic limbs express a cluster of genes associated with cell proliferation and metabolism first, followed by gene clusters associated with patterning and morphogenesis.

In addition to investigating the expression dynamics of gene sets defined by clustering and GO term enrichment, we took a more targeted approach, focussing on the *Parhyale* orthologues of genes that are involved in immune responses, cell proliferation, limb patterning and cell differentiation (Figure 6, complete list in Supplementary Materials). The genes associated with immune responses, cell proliferation and patterning were selected based on published information (particularly from *Drosophila*; http://flybase.org/); genes associated with differentiated neurons and muscles were selected from *Parhyale* single-cell transcriptomic data (A. Almazan, M. Paris and M. Averof, unpublished data).

**Figure 6.**
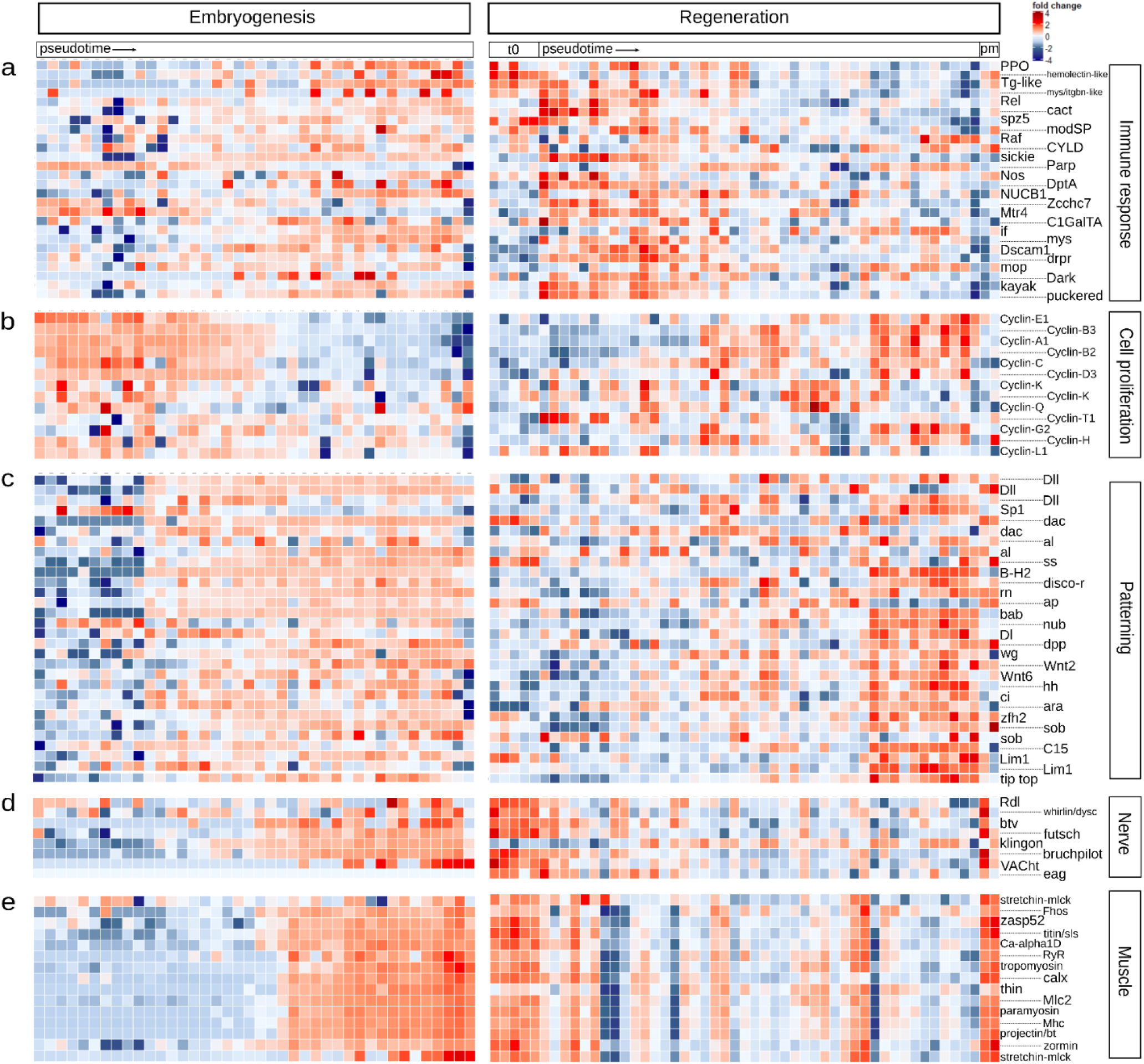
Heatmaps comparing expression in embryogenesis (E values, left) and regeneration (R values, right) for a selection of gene-sets: (**a**) immune response, (**b**) cell proliferation, (**c**) patterning, (**d**) nervous system, (**e**) muscle markers. t0: non-regenerating samples; pm: post-molt samples. *Drosophila* gene names are indicated for a selection of *Parhyale* orthologs. Samples were ordered according to hours post fertilization for the embryogenesis dataset and according to pseudotime for the regeneration dataset.

The comparison of transcriptional profiles for immune system-related genes (Figure 6a) shows that they are markedly upregulated during the early phases of regeneration dataset, following wounding. The same genes are expressed only in the later phases of embryonic development, possibly connected to the differentiation of circulating haemocytes.

The expression profiles of M and G1/S phase cyclins indicate that cell proliferation occurs mainly during the early phase of limb embryogenesis, and the mid-late phases of regeneration (Figure 6b). This is consistent with data from live imaging (Alwes et al. 2016) and with the GO enrichment analysis (Figure 5c,d).

Limb patterning genes were predominantly expressed in the mid-late phases of embryonic leg development (from 120 hpf onward), after the expression of genes associated with cell proliferation (Figure 6c). During leg regeneration, this set of genes is predominantly expressed during late phases, overlapping with the expression profiles of genes associated with cell proliferation (Figure 6c).

Genes associated with differentiated muscles and nerve cells are strongly expressed during the late phases of leg development in the embryo, and in fully differentiated adult limbs before and after regeneration (Figure 6d,e). They are strongly downregulated during the time course of leg regeneration, possibly reflecting neuron and muscle de-differentiation during these stages.

## CONCLUSIONS

Comparing the transcriptional profiles of developing and regenerating legs has allowed us to probe whether the process of leg regeneration recapitulates parts of embryonic development, in terms of transcriptional dynamics, on a global, genome-wide scale.

First, we find that the transcriptional dynamics of leg development are stereotypic and highly reproducible across individuals (Figure 1). The developmental stage of a limb can be predicted from the transcriptome to within ∼10% of developmental time (Figure 1c). In contrast, the transcriptional dynamics of leg regeneration are embedded within a high level of individual variation (Figure 2), which is largely driven by the molting cycle (Figure 3b). This is consistent with the fact that, unlike development which occurs in the relatively stable environment of the egg, regeneration takes place in the context of the complex physiology of the adult.

By removing the effects of individual variation, through Bayesian modelling, we are able to recover the transcriptional dynamics of leg regeneration (Figure 4). Like the dynamics of leg development in the embryo, these reveal distinct phases of gene expression that unfold during regeneration, with most variation occurring during early-mid regenerative stages (Figure 4c); this is consistent with variation in cell dynamics observed by live imaging (Alwes et al. 2016). Using GO term enrichment analysis we can assign putative gene functions to each of these phases. This analysis reveals distinct phases for wound healing, metabolic reprogramming (during a period previously described as a phase of quiescence, Alwes et al. 2016), cell proliferation and morphogenesis, and finally cell differentiation (Figure 5c and 6).

We have tried to relate these phases of leg regeneration to the time course of leg development, using global comparisons of gene expression (Figure 5a,b), comparisons based on clusters of co-expressed genes identified for development and regeneration (Figure 5c-g) and by comparing the transcriptional dynamics of genes involved in cell proliferation, patterning and cell differentiation (Figure 6b-e). While we observe that the same processes and overlapping gene sets are implicated in both leg development and regeneration, we find that the temporal order in which they are deployed is not the same. This is true not only for phases and processes that are likely to be unique to regeneration – e.g. wound healing, immune/stress responses, metabolic reprogramming – but also to processes like cell proliferation, patterning and morphogenesis, which we expect to be shared between development and regeneration.

We conclude that the time course of leg regeneration is not collinear with that of leg development.

## METHODS

### RNAseq design and sequencing

#### Embryonic dataset

*Parhyale* females of the Chicago-F inbred line (Kao et al. 2016) were collected after fertilization, and their embryos removed about 3 dpf. Each brood was kept separately, in a temperature-controlled incubator (set at 27°C), in sterile 6-well plates (Costar #3516), in filtered artificial seawater (FASW) with antibiotic. Embryos were staged 3-4 days post fertilization. The initial limb bud will give rise to the limb segments (dactylus-propodus-carpus-merus-ischium-basis), plus the coxal plates and the gills. In order to collect the distal-most part of the leg while coping with the progressive specification of regions along the proximal-distal axis of the limb, two different distal leg datasets were collected: a dataset of late samples (A series) comprising the carpus-propodus-dactylus region (144-192 hpf, 21 samples, Figure 1a) and a series of earlier samples (AB series) for which the merus and the rest of the leg are not differentially specified yet and comprising the prospective merus-carpus-propodus-dactylus regions (120-138 hpf, 9 samples). Individual full T4 limb samples were also collected every 6 hours for a broader time window, from 96 hpf to 192 hpf (F series, 40 samples in total). At least 2-3 samples were collected for each time point and each condition. Embryos were dissected in the lid of a 5 ml Protein LoBind Tubes (Eppendorf #*0030108302*), in 1% BSA in FASW (80 ul). The eggshell was removed with fine forceps (Fine Science Tools #11254-20), and limbs were dissected with borosilicate needles (pulled capillaries; Sutter #B100-50-15). Samples were transferred in 100 ul of ice-cold lysis solution (Agilent Absolutely RNA Nanoprep kit #400753), homogenized though brief pipetting, and flash-frozen in liquid nitrogen. RNAextraction was performed with the Agilent Absolutely RNA Nanoprep kit (#400753), following manufacturer’s instructions, and eluted in 10 ul of elution buffer. RNA extraction was randomized. As the concentrations of RNA extracts were too low to be directly quantified, they were treated as follow: 9 ul from each samples were directly used for cDNA amplification (15 cycles of LD PCR, using the SMART-Seq v4 Ultra Low Input RNA kit for sequencing (Takara Bio, #634898) and the SeqAmp DNA Polymerase (Takara Bio #638509), 1 ul of cDNA was then used for Qubit quantification (4.0 HS DNA), providing concentrations in the range of 0.2-0-7 ng/ul. Libraries were synthesized from 1 ng of cDNA, using the Nextera XT DNA Library Preparation Kit (Illumina #FC-131-1096; with Dual indexing strategy, i7 and i5), and with an optimised protocol that included an accelerated cooling step on ice after the 55°C step. Quantification and validation of libraries were done with both Qubit 4.0 (HS DNA kit, Thermofisher) and Tapestation D5000 equipment using the D5000 ScreenTape System (#5067-5588 and #5067-5589, Agilent). QC libraries were normalized and then loaded into an Illumina NextSeq 500 sequencing system using NextSeq 500 High Output Kit with 76 bp single-end sequencing, according to the manufacturer’s instructions (Illumina, San Diego, CA, USA).

Details about the sequenced samples are provided in Supplementary materials.

#### Regeneration dataset

Wild-type (Chicago-F line) and Distal^DsRed^ transgenic (gene trap where DsRed fluorescent protein is cytoplasmically expressed in the distal region of all appendages, Kontarakis et al. 2011) animals were selected as described above, and individually kept in homogeneous conditions for three months prior to experiment. In order to test for the inter-and intra-individual variability of the regenerative process, both T4 limbs of each animal were amputated simultaneously, proximally to the carpus/propodus joint. Samples from the Chicago-F line were harvested pre- and post regeneration, and every 12 hour, from 0 to 120 hpa; samples from the Distal^DsRed^ line were collected when the DsRed signal became visible (around 150 hpa). Samples from the same individual were collected at the same moment. Due to the observed variability in the regenerative sequence, samples were processed and sequenced individually, as follows: *i)* from each animal, either both or only one T4 limb was harvested, *ii)* from each limb, two fragments were isolated, one including the regenerating podomere(s) (A fragment, localized on the carpus - for consistency of the input material, we always collected the same region, also from non-regenerating limbs) and one its proximal control podomere (B fragment, localized on the distal part of the merus from the same limb). Five paired samples were collected per time point, with the exception of Distal^DsRed^ line and post-molt samples (2 samples each). In total 60 regenerating and 60 control samples were collected; a scheme is provided in Figure 2a, and more details about the samples are provided in Supplementary material. Limb fragments were immediately transferred in 1.5 ml LowBind tubes with 500 ul of ice-cold lysis buffer (Reliaprep RNA Tissue MiniPrep System, Promega #Z6111), vortexed, then transferred to a sterile multiwell plate for manual disruption of cuticle with a clean surgical blade. The sample was then re-transferred to the tube, vortexed, and frozen in liquid nitrogen, for then being stored at -80C until further processing. RNA extraction was randomized, avoiding to process at the same time related samples. RNA was extracted with the Reliaprep RNA Tissue MiniPrep System (Promega #Z6111), and eluted in 15 ul of nuclease-free water (Invitrogen #AM9937). RNA quality was assessed with TapeStation D5000 (Agilent RNA ScreenTape High Sensitivity system #5067-5579, #5067-5580. #5067-5581), and 1 ng of RNA was used for cDNA synthesis (SMART-Seq v4 Ultra Low Input RNA kit – Takara Bio: #634898). Verification of library quality and sequencing were done as for the embryonic dataset (see above). Details about the sequenced samples are provided in Supplementary material.

#### Molting dataset

Chicago-F animals, selected as previously described, were individually kept and monitored for 3 months, in order to determine their molting cycles. The data was used to estimate collection times: 5 days before molt, 1-2-3 days before molt, 1 or 2 days after molt, 9-10 days after molt. One intact, full T4 limb was harvested per animal, and RNA was extracted as described above. A total of 20 limbs deriving from animals at different stages of their molting cycle was sequenced. Library preparation and sequencing were done as described above. Details about the sequenced samples are provided in Supplementary material.

#### Adult full limb dataset

In order to help build a new reference transcriptome (see later), two additional full T4 limb samples from non-regenerating Chicago-F males were collected and processed as described above. Details about the sequenced samples are provided in Supplementary material.

### Reference transcriptome assembly

We used the *Parhyale hawaiensis* genome assembly Phaw_5.0 (https://www.ncbi.nlm.nih.gov/assembly/GCA_001587735.2/) to which the following modification was done: 19 poorly assembled mitochondrial scaffolds were removed from Phaw_5.0 and replaced with the publicly available complete mitochondrial genome (RefSeq: NC_039402.1).

Gene models were obtained as follows: reads obtained from the regeneration, embryogenesis and adult full limbs datasets (for a total of 2,656,106,490 reads) were mapped to the modified *Parhyale* genome described above, using hisat2 v2.1.0 with the default parameters plus “--dta” to increase the threshold of anchor length. Then gene models were first inferred based on the publicly available annotation (https://www.ncbi.nlm.nih.gov/assembly/GCA_001587735.2/) using stringtie v2.0.3 with stringent conditions: “-c 10 -j 10 -t -f 0.20”. This annotation was further merged with the publicly available one using “stringtie --merge”. All transcripts with length below 500 bp were removed. Despite those stringent conditions, we noticed some over-annotation with overlapping genes. We removed genes that overlapped by more than 50% of their transcript length with another gene that is longer and on the other strand if they didn’t show any good blast hit (score < 50 or e.value > 0.01). We also noticed that some genes were wrongly split in multiple genes in the annotation. Those putatively split genes were identified based on blastn to assembled sequences from *Parhyale* Iso-Seq (see below) or blastp to the related species *Drosophila melanogaster* (Uniprotprotein set UP000000803_7227) and *Hyalella azteca* (predicted protein set from the i5k project: hyaazt_OGSv1.2_pep). The procedure for flagging putatively split genes was as follows: two nearby *Parhyale* genes (distance below 1Mb) were considered as belonging to the same genes if they shared the same Blast hit with another gene in the Iso-seq dataset, *Drosophila* proteins or *Hyalella* proteins, meeting the following conditions: *(i)* high percent sequence identity (over 95% for the Iso-seq reads, 40% for *Drosophila* and 60% for *Hyalella*); *(ii)* overlap on long enough stretches (over 300 nt for the Iso-seq or 50 aa for *Drosophila* and *Hyalella*); *(iii)* at coherent positions (e.g. the *Parhyale* gene located in 5’ blasts to the 5’ end of a *Drosophila* protein while the nearby *Parhyale* gene located in 3’ matches the 3’ end of the same *Drosophila* protein). The merging of split genes was not included in the gtf file because it was not based on detection of splice variants. Merging was taken into account after read mapping by simply adding up counts or tpm of the split genes.

The annotation was further manually curated, dealing with a handful of duplicated transcripts; a manually curated Ubx sequence and an RFP sequence were added to the fasta file of transcript sequences used for further analyses.

The final gtf file contains 54718 genes and the final list used for further analysis includes 52759 genes (Supplementary Materials).

#### PacBio long-read sequencing (Iso-seq)

Adults and juveniles of the Chicago-F line were provided by the Patel Lab at the University of Chicago Marine Biological Laboratory (October 2019). Fifteen animals were combined, seawater removed, and high quality total RNA extracted (RIN 8.5) following the Qiagen RNeasy kit protocol (catalog number 74004) and using rotor-stator homogenization. cDNA synthesis and a PacBio sequencing library (chemistry 2.0) was prepared according to kit protocols for subsequent PacBio Sequel II sequencing. Reads were processed using the SMRTLink IsoSeq3 pipeline (version 10) and produced a final transcriptome of 24,962 transcripts. The transcriptome was assessed using EvidentialGene (version 19may14; 1000 longest “okay” transcripts (21,776 total): Average length 1,378; median length 1,255; shortest 972; longest 3,649 amino acids) and BUSCO (version 4.0.6), with BUSCO evaluation using the BUSCO Metazoa database (version metazoa_odb10; BUSCO Metazoa transcriptome completeness: 59.7% Complete; 57.9% Complete single copy; 1.8% Complete duplicated).

### Orthology inference and GO annotation

Orthology search against *Homo sapiens, Drosophila melanogaster, Ambystoma mexicanum*, and *Hyalella atzeca* was done using BLASTP, and Best Reciprocal Blast Hits were selected. Annotation was complemented with eggNOG 5.0 (Huerta-Cepas et al. 2019), which assigned a GO term to 15261 sequences.

### Measurements of gene abundance

For both embryonic and adult datasets, reads were mapped to the 54718 genes of our annotation (plus RFP), using kallisto v. 0.42.5 (Bray et al. 2016) with the following parameters: –single -l 350 -s 50 -b 100. The output was a matrix of transcript-level abundances. Gene abundances were obtained by simply summing over all the transcripts of a given gene. Given that multiple sequences were flagged as being fragments of the same gene, we further added up a posteriori the expression values of those putative split genes.

### Analyses of the sequenced datasets

Genes in the embryonic dataset or in the regeneration dataset, for which fewer than 20 reads mapped, were separately removed from the analysis: 432112 and 43968 genes were kept for the embryogenesis and regeneration datasets, respectively.

#### Normalization

Count and tpm matrices were first quantile-normalized (limma R package, v. 3.48.0; Ritchie et al. 2015), then log transformed (log(x+1)). The JAGS-transformed datasets were only log2 transformed.

#### Principal Component Analysis, heatmap

PCAs were performed using the R function prcomp (parameter scale set to TRUE), and the 10000 most variable genes. Heatmaps were plotted with the ComplexHeatmap package (v. 2.8.0; Gu et al. 2016), which required the additional dendextend (v. 1.15.1) and circlize packages (v. 0.4.13; Gu et al. 2014).

#### Differential expression analysis

We used the R package DESeq2 (Love et al. 2014) on raw counts for identifying genes differentially expressed during embryogenesis, separately in the full limb and the A/AB series (using hpf as the explanatory variable). Genes with a p.adj < 0.001 were selected. In order to determine the list of molting-related genes (which we used at different steps to asses the removal of the physiological signal), we also applied DESeq2 to identify the genes differentially expressed between the different time windows of the molting dataset (p.adj < 0.001): the union of significantly differentially expressed genes from all pair-wise comparisons was selected as our list of diagnostic molting genes.

#### Identification of co-expressed gene sets

We applied a soft clustering approach, using the R package mFuzz (v. 2.52.0; Kumar and Futschik 2007), setting sd = 0.2, and membership = 0.8 for calculating the eset object. The optimal number of clusters was estimated from the inflexion points of the Dmin function (iterations = 100): we identified 9 or 25 clusters for the R dataset, 4 or 33 for the E, and 8 for the molting one (see Supplementary materials). In order to plot expression dynamics, we extracted the cluster centroids values, which were used to build an input matrix for the heatmap (see Figures 3c, 5c-d).

#### Comparison of clusters

Fold enrichment scores were computed as follows: we took an approach inspired from the one described in Kowarsky et al. (2021) where they use an hypergeometric distribution to estimate an enrichment score between gene sets. The enrichment fold was calculated as the ratio of the number of genes that overlapped between two clusters over the size of an overlap that would occur with random sets of genes of the same size. Results were plotted as a heatmap, where values between 0 and 1 indicate no enrichment, while values >1 indicate enrichment.

The clustree package (v. 0.4.3; Zappia and Oshlack 2018) was used to assess the correspondence between the higher/lower numbers of clusters. The chord diagram was plotted with the circlize package (v. 0.4.13; Gu et al. 2014).

#### GO enrichment analysis

Enriched GO terms were identified using the packages clusterProfiler (v. 4.0.0; Yu et al. 2012), org.Dm.eg.db (v. 3.13.0; Carlson 2019) and enrichplot (v. 1.12.0; Yu 2021). The previously identified *Drosophila* orthologs of *Parhyale* genes (see “Orthology inference and GO annotation” section) were used as background for enrichment, additionally we set: pvalueCutoff = 0.05 and qvalueCutoff = 0.10. The list of GO terms was further trimmed for display (Figure 5c-d), by using the Revigo algorithm (http://revigo.irb.hr/ ; Supek et al. 2011). We set the following parameters: allowed similarity = small (0.5), semantic similarity measure = SimRel.

### Bayesian modeling of regenerating and embryonic datasets

Computations were performed using JAGS via the R package rjags (Plummer, 2019, see Supplementary Material). In the model described in Figure 4a, for each gene and each A and B sample, expression values (counts) are Poisson distributed from means uA and uB, respectively. uA values are the product of uB (gene and individual specific) and R (gene and sample specific). The log of R is Normally distributed (mean of 0 and variance sigma_R). uB is log-normal distributed (mean of ulog_B and variance sigma_B). sigma_R and sigma_B are uniformly distributed between 1 and 10 while ulog_B is uniformly distributed between -5 and 10.

Due to the nature of the embryonic samples, we did not have an equivalent to the regeneration B series. We decided to use the B samples from the four pre-amputation individuals, chosen because they did not show any molting signal (Figures 2c and 3b) nor were affected by regeneration. We hypothesized that the embryogenesis signal would not be present in the pre-amputation individuals that have fully formed legs in a steady-state condition so it would be preserved in the output of the model. An “embryogenesis factor” E was modelled as representing the fold change between the embryogenesis samples and a control expression level in the adult t0 B samples. We used the same priors as for the regeneration dataset (described above, Figure 4a).

Three to four independent MCMC chains were run in parallel for each model. For each chain, 21,000 samples were produced. The first 1,000 were considered as burn-in phase and discarded. To avoid autocorrelation, the remaining 20,000 samples were thinned by selecting one out of 100 samples, thus keeping 200 samples per chain. We checked the convergence by displaying MCMC chain traces and autocorrelation plots (R function autocorr.plot from the coda package) and by computing the Gelman and Rubin’s statistics as modified by Brooks and Gelman (1998) (R function gelman.diag from the coda package). For each parameter, its point estimate was defined as the median of its marginal posterior distribution.

### Predicting regeneration and developmental stages using RAPToR

In order to build predict stages of regeneration or development based on transcriptomic data, we used the R package RAPToR v1.1.4 (Real Age Prediction from Transcriptome staging on Reference), a recently developed tool to accurately predict individual samples’ developmental age from their gene expression profiles (https://github.com/LBMC/RAPToR).

For building RAPToR references we used the function ge_im with the formula formula = “X ∼ s(hpa, bs = ‘ts’) and parameter dim_red=“pca”. For the reference shown in Figure 1c, we used the top 20k most variable genes as calculated by DESeq2 on the raw counts and 32 PCs. To build the reference for Figure 2e, we used the top 20k most variable genes as calculated by DESeq2, excluding the genes with a low expression (median count below 10, with a final set of 10128 genes) on the raw counts and 4 PCs. To build the reference pseudotime for Figure 4c, we used the 20k top genes with the most variable R values also excluding the genes with a low expression (median count below 10), excluding the genes identified as differentially expressed in the molting RNAseq experiment (final set of 10531 genes) and only 4 PCs, to avoid overfitting (since we are directly using the training set). For the reference built for Figure 5b, we used the same gene list as for Figure 4c, further excluding genes not expressed in the embryogenesis dataset (for a total of 10152 genes) and 33 PCs.

### Graphics

The list of R packages used for making the figures are listed in the supplementary materials. Figures and illustrations were generated with the free software Inkscape (https://inkscape.org/).

## ACKNOWLEDGEMENTS

We thank Romain Bulteau for his help with RAPToR, Philippe Veber and Bastien Boussau for their help with rjargs, Nipam Patel for the *Parhyale* used for the Iso-seq experiment and the Salt Lake City sequencing facility for sequencing the Iso-seq libraries. We thank Enrique Arboleda for technical support, Pascale Roux and Laetitia Lebre for technical support on building the sequencing libraries. This work was funded by the European Research Council, under the European Union Horizon 2020 programme (project ERC-2015-AdG #694918). Alba Almazán was supported by the Marie Curie ITN programme ‘EvoCell’, under the European Union Horizon 2020 programme (project H2020-MSCA-ITN-2017 #766053).

